# Large-Scale Assessment of the Iranian population structure of Mitochondrial and Y-chromosome Haplogroups

**DOI:** 10.1101/2024.03.14.585067

**Authors:** Neda Mazaheri, Amin Ghahremani, Masoumeh Babazadeh, Damoun NashtaAli, Seyyed Abolfazl Motahari

## Abstract

The Iranian plateau, strategically positioned as a corridor for population diffusion across Eurasia, holds a pivotal role in elucidating the dynamics of human migrations originating from Africa around 60,000 years ago. Both prehistoric and historic movements of populations between Africa, Asia, and Europe may have been influenced by the unique geographical features of the Iranian plateau. Iran boasts ancient cultures and urban settlements predating some of the earliest civilizations, including the Neolithic revolution in neighboring Mesopotamia. Spanning from the Balkans and Egypt in the west to the Indus Valley in Pakistan and northern India in the southeast, the Iranian plateau encompasses a vast area characterized by incredible ethnocultural diversity. This region served as the origin for numerous mt-DNA/Y-DNA haplogroups that expanded to West Asia, Europe, Siberia, Central Asia, and South Asia. By examining both maternal and paternal haplogroups within the Iranian context, we aim to contribute to the broader narrative of human dispersals and elucidate the role those specific regions, such as the Iranian plateau, played in shaping the observed genetic diversity today. Due to the lack of comprehensive studies on mt-DNA /Y-DNA haplogroups in the Iranian population, our study sought to uncover the distribution of haplogroups among Iranian peoples using a large sample size. Our analysis focused on the frequency of ancestral haplogroups in Iran through the examination of large-scale whole-exome sequencing (WES) and SNP microarray data from 18,184 individuals. In our study, we observed 24 mt-DNA super haplogroups in the Iranian population, with the most common haplogroups belonging to West-Eurasian lineages U (20.73%), H (18.84%), J (12.10%), HV (9.22%), and T (8.98%), collectively comprising 69.70% of all Iranian samples. Notably, subclades J1 and U7 emerged as the two most frequent subclades, with frequencies of 11.24% and 7.30%, respectively. We also revealed the presence of 14 distinct Y-DNA haplogroups, with J, R, G, T, and Q emerging as the five predominant lineages. Notably, J2 (including J-L26) exhibited the highest frequency at 35.64%, followed by R1a at 14.68%. also, The detected mtDNA and Y-chromosome haplogroups were clustered into distinct groups that confirmed the heterogenicity of the Iranian population because of various factors including geographic or linguistic ethnic groups.

## Introduction

Progression of genomics technologies over the last few decades has sparked a significant revolution in the fields of population genetics and archaeology. The ability to decode the genetic information of individuals has played a pivotal role in addressing historical inquiries [1]. Genetic ancestry testing, a key component of this revolution, involves the examination of mitochondrial DNA (for tracing maternal haplogroups), Y chromosome DNA (for tracing paternal haplogroups), and autosomal DNA variants inherited from both parents. These genetic markers are routinely genotyped using DNA microarrays, and the resulting data is compared with the frequencies of these variants in reference populations sampled from around the world. Through this process, the geographic region with the highest frequency of a specific variant is considered the most likely origin of an ancestor who transmitted the variant to the person being tested [2]. Consequently, this testing methodology offers valuable insights into the ancestral histories of human populations, providing detailed information on migration patterns and distribution maps of populations worldwide [3]. Owing to its distinctive geostrategic location, the Persian Plateau has served as a crucial crossroads for human migrations spanning the African, Indian, and Eurasian plates since the emergence of modern humans from Africa. The presence of numerous prehistoric sites scattered across the Iranian plateau attests to the existence of ancient cultures and urban settlements that predates the earliest civilizations in the neighboring Mesopotamian region by several centuries [4]. The population of the Iranian plateau has undergone significant fluctuations due to migratory influxes throughout the past millennia. The inception of the first agricultural revolution in the Zagros, which radiated both westwards and eastwards, exerted profound influences on the demographic landscape of the surrounding regions of the Iranian plateau [1]. The Persian Plateau spans from the Caucasus mountains and the northern coast of the Black Sea in modern-day Georgia, Armenia, Azerbaijan, Turkey, and Russia in the north, to the Arabian Sea and the Persian Gulf in modern-day Iraq, Kuwait, Bahrain, Qatar, Oman, Yemen, and Saudi Arabia in the south. It extends from the Balkans and Egypt in the west, parts of West Asia as the base, the majority of Central Asia to the northeast, and parts of South Asia, including the Indus Valley in Pakistan and northern India, to the southeast [5, 6] encompassing this vast area is an incredible ethno-cultural diversity [7, 8]. Due to Iran’s strategic location on the Silk Road, connecting the Roman Empire and the Han Dynasty in China, it rapidly evolved into a hub of trade and commerce. Present-day Iran shares land borders with multiple nations: Iraq to the west, Turkey, Armenia, and Azerbaijan to the northwest, Turkmenistan to the northeast, and Afghanistan and Pakistan to the east. To the south, Iran is bordered by the Persian Gulf, serving as a maritime gateway between Iran and the eastern coastal region of the Arabian Peninsula. This area encompasses Kuwait, Saudi Arabia, Bahrain, Qatar, the UAE, and Oman. Notably, the Strait of Hormuz, the narrowest section of this waterway, separates southwestern Iran from the northeast tip of Oman by a mere 46.7 km of shallow water, punctuated by numerous tiny islands [9, 10, 11]. It is crucial to acknowledge certain geographic features in present-day Iran, such as the expansive deserts of Dasht-e Loot and Dasht-e Kavir, as well as the formidable Zagros and Alborz mountains. These features have served as significant barriers, profoundly influencing the genetic diversity of populations across this vast region (4). The continuous influx of populations with diverse origins and cultures has led to a remarkable amalgamation of various ethnic groups, each speaking a variety of languages within Iran. Consequently, a genetic evaluation of ancestral haplogroup distributions in Iran provides valuable insights into the historical migration routes and settlements of populations in this expansive and diverse area. This project represents the first large-sample-sized study on ancestral haplogroup analysis in the Iranian population, encompassing a majority of provinces and ethnic groups in Iran.

## Material and Method

### Sample collection & extraction

We collected samples from 18,184 unrelated individuals of Iranian descent for this study. The predominant sample type utilized in this study was blood, from which DNA extraction was conducted via the salting out method, and whole-exome sequencing samples were performed on 16933 samples. Additionally, 1251 saliva samples were collected using two buccal swabs and DNA extraction was performed using the salting out method.

### Sequencing and Analysis

Genotyping utilized the Infinium Global Screening Array-24 v3.0 kit (Illumina, San Diego, CA, USA), following the manufacturer’s protocol. Sample preparation and subsequent genotyping were conducted on an Illumina iScan system. 1251 samples used for MT-chromosome analysis while a subset comprising 760 samples underwent Y-chromosome analysis (male samples), targeting 778,784 DNA markers. Furthermore, for whole exome sequencing, the Agilent SureSelect-V7 kit was used for library preparation. Paired-end sequencing with coverage of minimum 100x was performed by the Illumina NovaSeq6000 sequencer machine. Reads were aligned to the human genome (UCSC GRCh37) using the Burrows-Wheeler Aligner (BWA). Detection of functional variants from high-throughput sequencing data was carried out using the Genome Analysis Toolkit (GATK) software. Note that, mitochondrial variants are off-target of the Agilent SureSecelct-V7 kit, but was targeted with good quality. For MT-chromosome analysis, within WES and SNP chip, we select an on the average 32 SNPs per sample and exactly 384 SNPs for further analysis, respectively. Haplogrep, a tool developed by Institute of Genetic Epidemiology (Medical University of Innsbruck), facilitated MT-chromosome haplogroup identification for both data types. For Y-chromosome analysis, within both WES and SNP chip data we select 2,302 SNPs (from on the average 3,567 SNPs per sample) were retained for further analysis. We use all variants which were cataloged in the international Y tree maintained by the International Society of Genetic Genealogy (ISOGG, 2019) for haplogroup inference. For genotyping data, Yhaplo are used for Y-chromosome haplogroup identification, employing a BFS algorithm. While, Yleaf was employed for bam file of the WES data, which demonstrating superior performance compared to benchmarked tools in whole exome sequencing (WES) data, successfully classifying samples [26].

### PCA and UMAP analysis

To assess the extent to which the defined haplogroups mt-DNA/Y-DNA diversity within the population, we performed unsupervised clustering of the subjects and assessed the consistency with haplogroup assignments. We employed two unsupervised classification approaches: principal component analysis (PCA) and uniform manifold approximation and projection for dimensionality reduction (UMAP). In the context of genetic data, UMAP identifies the nearest genetic neighbors for each individual and groups closely-related individuals together. UMAP has emerged as a widely used tool for studying population structure in humans and other species, often complementing existing methods. We utilized the silhouette method as a metric to identify the optimal parameters UMAP to discover the most informative representation of super haplogroups.

### Result and Discussion

#### mt-DNA haplogroup frequency

In this study, we observed 24 mt-DNA super haplogroups within the Iranian population. Our analysis revealed that the most prevalent mt-DNA haplogroups were U (20.73%), H (18.8%), J (12.10%), HV (9.22%), and T (8.98%) (Fig. 1). Among the mt-DNA sub-haplogroups, J1 (11.24%), U7 (7.3%), and T2 (5.1%) exhibited the highest frequencies in the Iranian population. Detailed frequencies of haplogroups and sub-haplogroups can be found in Fig. 1.

**Fig. 1.**
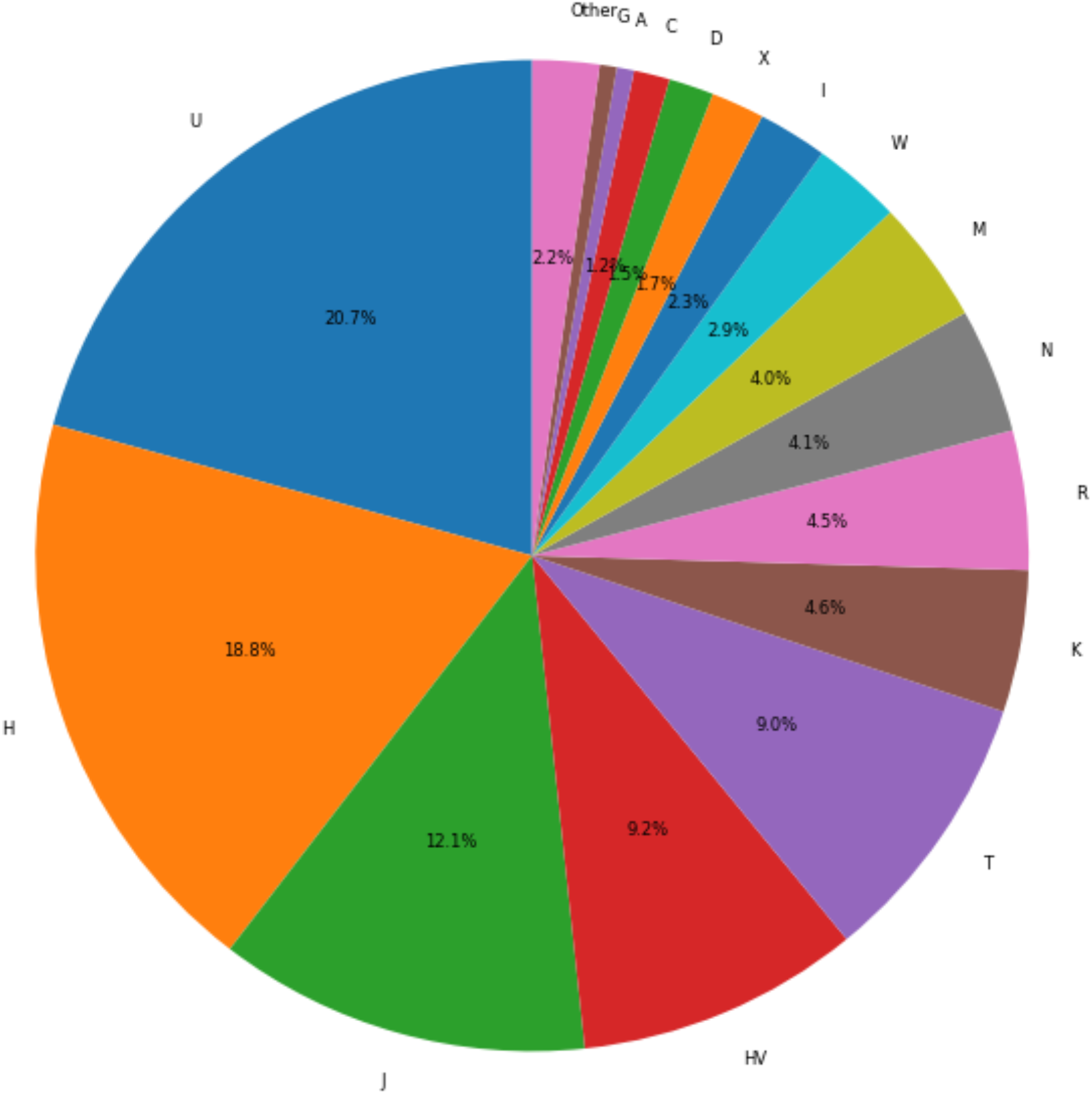

**Haplogroup U** emerged as the most prevalent maternal haplogroup in the Iranian population, accounting for 20.73% of the observed frequencies [figure1]. This haplogroup is recognized as specific to West Eurasia and is believed to have originated around 43,000 to 50,000 years ago, likely in West Asia, possibly in the Caucasus region [12,13]. Following Haplogroup U, U7 emerged as the second most common subclade at a frequency of 7.3%. Despite uncertainties surrounding its phylogenetic structure and precise ancestral origin, U7 is thought to have originated in the Black Sea area approximately 30,000 years ago [14]. The oldest ancient Iranian sample of U7 was recently detected in Southwest Iran, dating back approximately six thousand years (Kya) [15]. This haplogroup exhibits generally low population frequencies with restricted diversity across a wide geographic region, spanning from Europe to South Asia, including around 4% in the Near East and peaking at 10% in Iranians. Previous reports have also highlighted the presence of the U7 haplogroup at significant levels in northern Iran [16,17]. Notably, our analysis revealed that U7a (6.30%) and U7b (0.55%) are the predominant subclades of U7 observed in the Iranian population.

**Haplogroup H** emerged as the second most common mt-DNA haplogroup in the Iranian population, with a frequency of 18.84%. It is believed to have originated in Southwest Asia near present-day Syria approximately 25,000 years ago [18]. Initially, it expanded into the northern Near East and Southern Caucasus before diffusing from Iberia to Europe prior to the Last Glacial Maximum [19]. The H macro-haplogroup, along with U, is considered a predominant lineage within Europe, constituting around 40% of all maternal lineages, and in the Near East, including 20% in the Caucasus and 17% in Iran [20]. Previous studies have consistently identified H as one of the most frequent maternal haplogroups in Iran [9, 21, 22, 23], consistent with our findings. Within our Iranian samples, we observed H13 (including H13a2a) at a frequency of 2.60%, followed by H14 (2.08%), H11, H2 (1.30%), and H5 (1.06%) as the most prevalent subclades of haplogroup H, respectively.

**Haplogroup J**, with a frequency of 12.10%, emerged as the third most prevalent mt-DNA haplogroup in our Iranian samples, following U and H. This finding is consistent with previous studies that have highlighted the substantial presence of haplogroup J as a major maternal component in Iran [9, 16, 17, 21]. Haplogroup J is believed to have originated approximately 45,000 years ago, likely in West Asia, specifically the Near East or Caucasus regions. Interestingly, recent research by Kristjansson et al. revealed that mitochondrial DNA haplogroup J is also the third most frequent haplogroup in modern-day Scandinavia, despite not originating there [24]. Within our study, the majority of haplogroup J1 was represented by J1a (frequency of 11.24%), with J1b (including J1b1b1) being the more prevalent subclade at a frequency of 7.52%. Notably, J1b subclades, particularly J1b1, exhibit higher frequencies among non-R1b populations in the Middle East, notably in the South Caucasus, Iran, and the Arabian Peninsula, albeit with distinct subclades. Other subclades of J1b are primarily restricted to the Middle East or the eastern Mediterranean.

**Haplogroup HV**, with a frequency of 9.22%, emerged as the fourth most common haplogroup in the Iranian population. This observation aligns with expectations, as Western Iran is considered a potential place of origin for HV. Furthermore, our findings confirm previous reports indicating that HV is among the more common maternal haplogroups in Iran [9, 16, 17, 21]. Within our study, the more predominant subclades of HV included HV2 (1.59%), HV14 (1.46%), and HV1 (1.26%).

**Haplogroup T** emerged as the fifth most common maternal haplogroup in the Iranian population, with a frequency of 8.98%. Within the T haplogroup, the T2 subclade was more prevalent, accounting for 5.17%, compared to T1 at 3.59%. Haplogroup T is believed to have originated in the Near East approximately 25,000 years ago and previously investigations in Iranian population (in small sized sample) were found this lineage at frequency of 2-3% [9,16,17]. Our finding confirmed that reports at a large sized sample of Iranian population.

In summary, our analysis revealed that maternal haplogroups U, H, J, HV, and T collectively account for over 69% of the maternal haplogroups observed in Iranian individuals. Conversely, macrohaplogroups K, R, N, and M were found at lower frequencies, ranging from approximately 4.6% to 4%. Additionally, haplogroups W and I were identified at frequencies of 2.9% and 2.3%, respectively, while haplogroups X (1.7%), D (1.4%), and C (1.16%) were classified as rare maternal haplogroups. Furthermore, very rare maternal haplogroups in the Iranian population, each comprising less than 3.3% of the population, include haplogroups A, G, F, B, V, Z, Y, P, and S, which are more commonly found in East/Southeast Asian, Indigenous Australian, and Oceania populations. Notably, haplogroup V, relatively rare in the global context, was found in around 4% of native Europeans [25].

### Y haplogroups frequency

As depicted in Fig. 2, the identified Y-DNA haplogroups in our study encompass 14 super haplogroups (J, R, E, G, T, Q, H, C, N, L, I, O, B, D, and K). Notably, the Iranian population exhibits remarkably high haplogroup diversity, and this study represents the first large sample-sized investigation into Y-DNA haplogroups within the Iranian population. Among the major Y-chromosomal haplogroups identified, J accounted for 46.20%, followed by R at 21.76%, E at 11.88%, G at 6.25%, T at 4.30%, and Q at 3.62%.

**Fig. 2.**
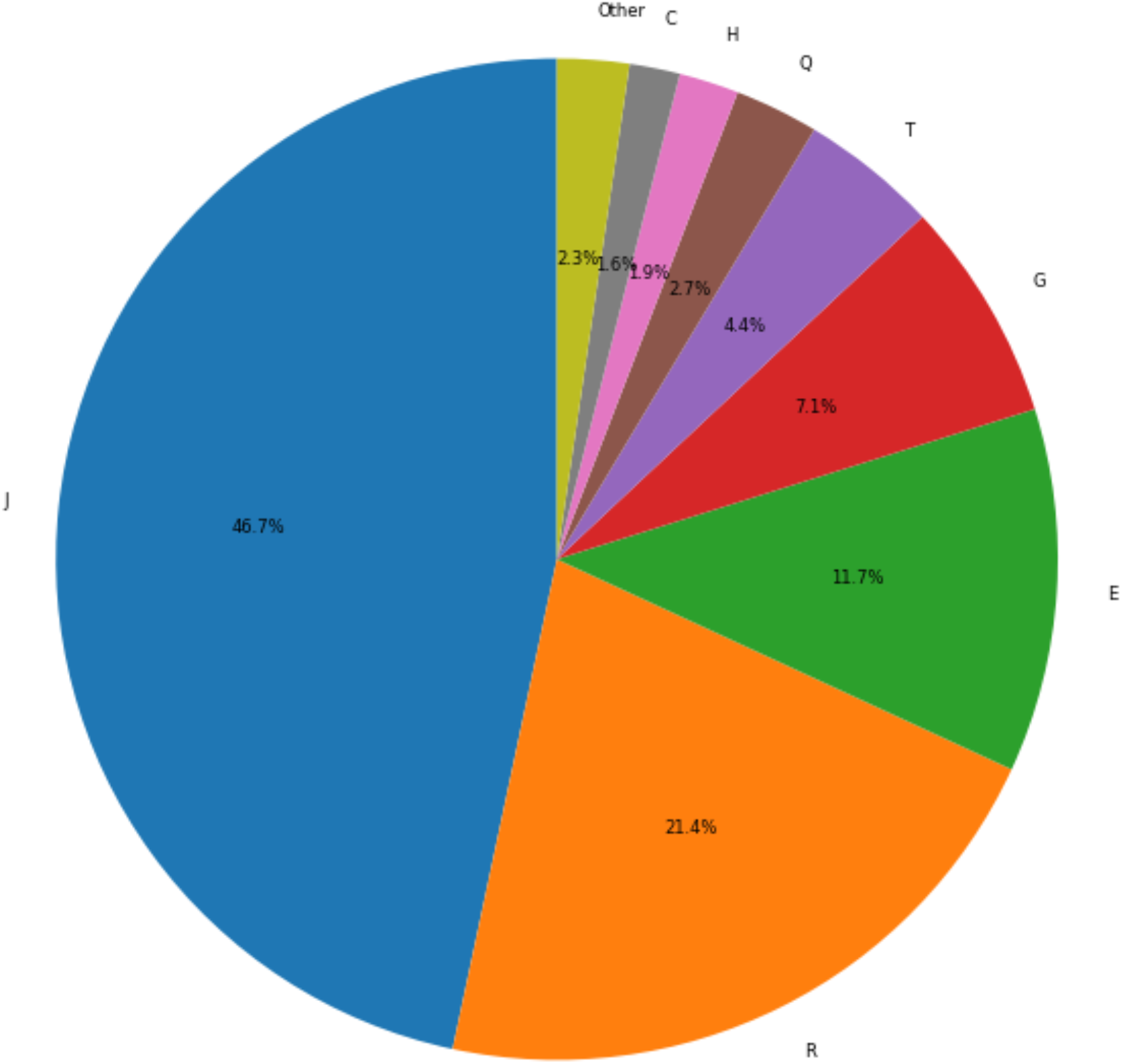

**Haplogroup J** emerges as the predominant Y-DNA haplogroup in the Iranian population, accounting for 46.20% of the samples, encompassing both main sub-clades, J2-M172 and J1-M267.

The J2-M172 subclade emerges as the most prevalent paternal haplogroup in the Iranian population, constituting 35.54% of the samples, as indicated in Fig. 2. This finding corroborates previous reports highlighting the common occurrence of J2 in the Iranian population [26,27]. The prevalence of J2 in Iran is anticipated given it believed origin in Western Iran [28, 29, 30] and its dispersion via Neolithic farmers from the Zagros mountains and northern Mesopotamia across the Iranian plateau to South Asia and Central Asia, as well as across the Caucasus to Russia (Volga-Ural). Subsequently, J2 is thought to have diffused with the advent of metallurgy from the southern Caucasus toward northern Mesopotamia and the Levant, and later propagated through Anatolia and the Eastern Mediterranean with the rise of early civilizations during the Late Bronze Age and the Early Iron Age [31]. It is noteworthy that among the observed subclades of J2-M172, the most common include J-L26 (10.54%) and J-M47 (4.47%) (Fig. 2). Among our Iranian samples, J-L26 emerged as the most highly frequent Y-DNA subclade, representing a novel finding as there has been no previous data reported on the frequencies of J2 haplogroup subclades in Iranian samples. Following J-L26, J-M47 was the second most prevalent subclade. It is known that J-M47 branches exhibit maximum phylogenetic diversity in regions such as Mesopotamia, the Persian Gulf, and Saudi Arabia, with major subclades being particularly prevalent in Armenia, Iran, South Asia, Anatolia, and Europe [32]. The frequency of J2-M47 in Iran has been reported in populations such as the Zoroastrians of Yazd, Mazandaran, Khuzestan, and Fars [27].

J1-M267 is believed to have evolved approximately 20,000 years ago, with its likely origins spanning northwestern Iran, the Caucasus, the Armenian Highland, and northern Mesopotamia. Presently, it is found at higher frequencies in the Arabian Peninsula and Western Asian countries, including Yemen, Qatar, Iraq, Saudi Arabia, and Palestine. Notably, this haplogroup is primarily found in Arab countries, while in Europe, it is relatively rare. Outside the Asian continent, J1-M267 is found in regions such as Sudan, parts of the Caucasus, Ethiopia, and North Africa, with high levels also observed in Jewish populations. In Iran, J1 has been reported to be present in frequencies below 10%, primarily found in regions such as Fars, Zoroastrians from Yazd, Gilan, Assyrians from Azerbaijan, and Khuzestan. Specifically, J1-Page08 is more common in populations residing below the Dasht-e Kevir and Dasht-e Lut desert area (approximately latitude 30°N), reaching its highest frequency among Arab populations in Khuzestan, bordering southern Iraq [27]. In this report, the frequency of J1 in the Iranian population was found to be 10.66%, ranking as the fourth most common among all Y-DNA haplogroups, with J-P58 observed as the predominant subclade at a frequency of 4.83%. It is estimated that the J1-P58 variety evolved around 9,000 to 10,000 BCE in the Fertile Crescent, beginning its expansion from the southern Levant (Israel, Palestine, Jordan) across the Arabian Peninsula during the Bronze Age. Its migration path is believed to have originated from eastern Anatolia and the southern Caucasus, traversing through the Taurus and Zagros mountains, and eventually reaching Mesopotamia and the Arabian Peninsula at the end of the last Ice Age (12,000 years ago), facilitated by the seasonal migrations of goat and sheep pastoralists. Presently, this subclade constitutes the majority of male lineages in the population of the Arabian Peninsula but is less common in the Caucasus, Anatolia, and Europe [33].

#### R haplogroup

In previous studies, haplogroup R in the Iranian population has been predominantly composed of two R1 sub-lineages, R1a-M198 and R1b-M269, while R2-M124 was found at a low frequency (less than 3%) [27]. In our project, as depicted in Fig. 2, haplogroup R emerges as the second most common paternal haplogroup in the Iranian sample, with a frequency of 21.76%. Our results indicate that R1a is the most frequent subclade within the R haplogroup, with a frequency of 14.48%, followed by R1b at 4.26%, and R2 observed at a very low frequency of 1.5%. Evidence suggests that R1a likely originated in Central Asia or southern Russia/Siberia and, along with R1b, expanded to eastern Europe via Iran and the Caucasus [34]. R1b is believed to have first developed somewhere in northern Iran or southern Central Asia before diffusing to the Fertile Crescent for herding. Outside Western Europe, R1b is found in regions such as the Ural Mountains of Russia, the Caucasus, Turkmenistan, and parts of Africa. Houshmand et al. revealed that R1b haplogroup frequency reaches 4.3% among Iranian sub-populations, including Assyrians, Armenians, and those in Lorestan [27]. Finally, our findings confirm the rarity of R2 frequency in Iran, consistent with previous reports.

#### E haplogroup

Previous reports have indicated that haplogroup E in Iran is predominantly composed of E1 sub-lineages, particularly prevalent in regions such as Kurdistan, Lorestan, and among the Zoroastrians of Yazd [35]. In our report, haplogroup E emerged as the second most common Y-DNA haplogroup, with a frequency of 11.88% (Fig. 2). The majority of this haplogroup belongs to the E-V13 subclade (11.75%), a branch that originated approximately 22,500 years ago in the Middle East/Western Asia and subsequently diffused into the Balkans around 4500 years ago [36]. It is noteworthy that a small percentage of the population (0.13%, 7 out of 5131 individuals) belongs to the E2 subclade. E2 is an African-specific haplogroup, and Houshmand et al. reported three individuals of E2 in South-East Iran (Hormozgan and Sistan Baluchestan) [27].

#### Haplogroup G

emerges as the fourth most common Y-DNA haplogroup in the Iranian population, with a frequency of 6.25% (Fig. 2). Within this haplogroup, G2a is the most frequent sub-clade, accounting for 5.72% of the total sample (Fig. 2). Previous studies have described G2a as a more common sub-lineage of the G Y-DNA haplogroup in the Iranian population, with incidences ranging from 0% in Sistan Baluchestan to high levels in the Arabs of Khuzestan [27, 37, 38]. It has been suggested that the G2a subclade may have developed in West Asia, possibly in Anatolia/Iran, and expanded with the development of agriculture to the Caucasus and Europe. This lineage is now characterized by high frequencies in the Lebanese and Jewish populations. Additionally, G2a is found at low frequencies in regions such as the Arabian Peninsula, Syria, Iraq, Iran, Afghanistan, and Pakistan [38]. It is worth noting that G2b and G1a were also observed at low frequencies in our Iranian samples, at 0.05% and 0.44%, respectively (Fig. 2).

#### Haplogroup T

The T-chrY haplogroup predominantly represented by the T1a sub-clade, was found with a frequency of 4.30% in the Iranian sample (Fig. 2). T1a is believed to have originated during the late glacial period, approximately between 25,000 and 15,000 years ago, possibly in the Iranian Plateau, and is currently more common in Mediterranean Europe [39]. In Iran, the T-YDNA haplogroup is found at high frequencies among various populations, including Zoroastrians from Kerman, Bakhtiaris, Assyrians from Azerbaijan, Abudhabians, Armenians from Historical Southwestern Armenia, and Druzes from Galilee, peaking at up to 10%. It also reaches frequencies of around 4% of the population around the Zagros Mountains and the Persian Gulf, as well as around the Taurus Mountains and the Levant basin [27, 40]. T-L906, with a frequency of 2.08%, is a more frequent sub-lineage of T1a in our Iranian samples (Fig. 2).

#### Haplogroup Q

is predominantly represented by the Q2a (Q-M378) subclade, which was found at a frequency of 3.62% in Iranian samples. Previous reports have indicated high frequencies of Q in northern Iran, including among populations such as Grugni, Turkmens of Golestan, Isfahan (Persian people), Khorasan (Persian people), Lorestan (Luristan, Lurs), Azerbaijan Gharbi, and Fars (Persian people), with frequencies gradually decreasing towards the southwest [27, 37, 41].

**Other haplogroups:** H (1.93%), C (1.62%), I (0.74%), L (0.63%), N (0.37%), O (0.33%), B (0.19%) and D (0.03%) Y-DNA haplogroups were occurred at lower frequency (lower than 5.8% of totally paternal lineages) in the Iranian population and over than 94.12 % of paternal haplogroups was belongs to J, R, E, G, T and Q lineages (Fig. 2).

### mtDNA/ Y-chromosome variability

While the two-dimensional plot of PC1 and PC2 provided some insights, it proved challenging to fully capture the cluster classification information, including the distinct directions of different super haplogroups. Therefore, we opted for a three-dimensional plot comprising the top three PCs to better visualize and understand the underlying structure In Fig. 3 and Fig. 4, we present three-dimensional PCA, UMAP with the best parameters in two-dimensions and three-dimensions for MT-DNA and Y-DNA, respectively. In Fig 3., we see nine clusters of super-haplogroups within MT-DNA data of Iranian with PCA. Also, ten clusters of super-haplogroups can be observed with UMAP method. In Fig 4., we also see eight clusters of super-haplogroups within Y-DNA data of Iranian in both PCA and UMAP methods. These clusters may shed light on the ethnical or geographical heterogenicity of the Iranian peoples. Ethic /regional grouping of samples are needed for uncover the main factor which is corresponded to this clusters. These results also match with our ancestral information of Iranian population structure.

**Fig. 3.**
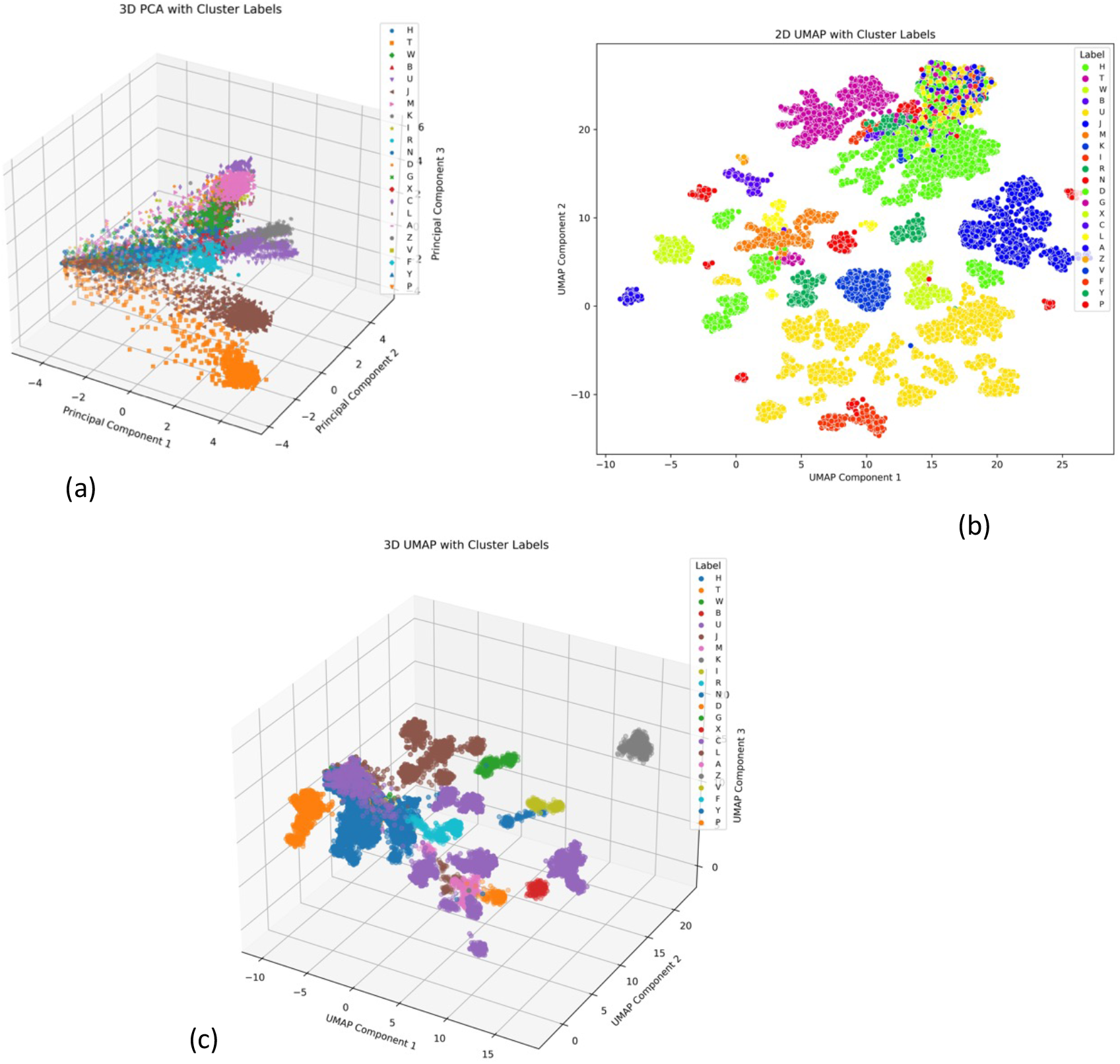
Clustering of MT-DNA data (both genotyping and WES samples) based on super-haplogroup, (a) three-dimentional PCA, (b) two-dimentional UMAP and (c) three-dimentional UMAP

**Fig. 4.**
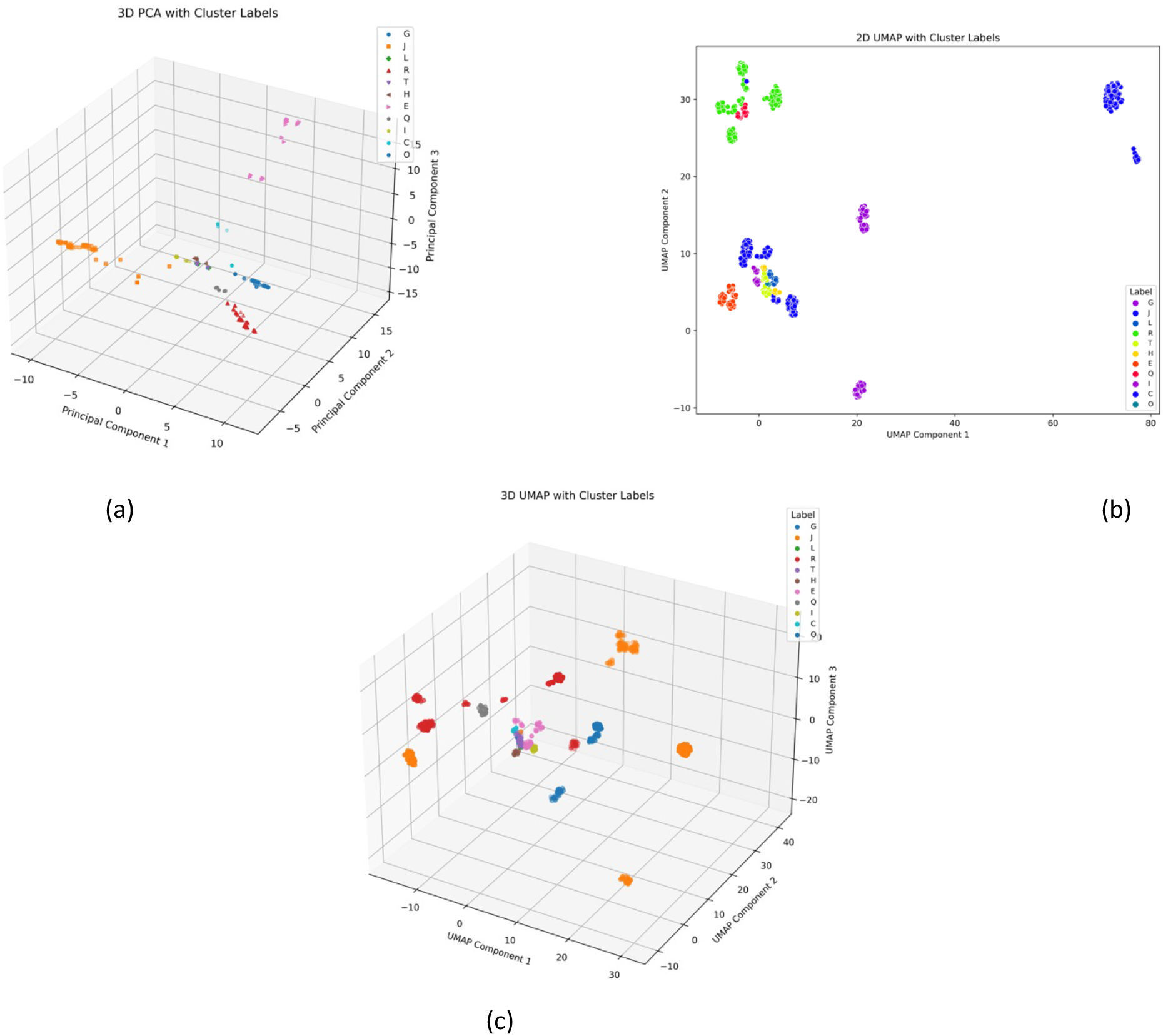
Clustering of genotyped Y-DNA genotyping data based on super-haplogroup, (a) three-dimentional PCA, (b) two-dimentional UMAP and (c) three-dimentional UMAP

## Conclusion

Given the lack of comprehensive studies on mitochondrial DNA (mt-DNA) and Y-DNA haplogroups in the Iranian population, our study aimed to elucidate the distribution of ancestral haplogroups among Iranian individuals using a large sample size. Specifically, we analyzed data from 18,184 individuals obtained from whole-exome sequencing (WES) and SNP microarrays. Our findings revealed 24 mt-DNA super haplogroups within the Iranian population, with the most prevalent haplogroups belonging to West-Eurasian lineages U, H, J, HV, and T, collectively accounting for 69.70% of all Iranian samples. Notably, the J1 and U7 subclades emerged as the two most frequent subclades, with frequencies of 11.24% and 7.30%, respectively. The analysis of Y-chromosome haplogroup frequencies revealed the presence of 14 distinct super haplogroups within the Iranian population, with the predominant haplogroups being J, R, G, T, and Q. Specifically, the J2 subclade, including J-L26, was the most prevalent at a frequency of 35.64%, followed by R1a (14.68%), E1 (11.61%), J1 (11.01%), G2 (6.71%), R1b (5.13%), and T1 (4.42%).

The heterogeneity of the Iranian mtDNA and Y-chromosome haplogroups in this project is well demonstrated by the PCA /UMAP analysis and detected distinct clusters of them. This complexity may have contributed to the various factors including geographic or linguistic ethnic groups and further exploration is needed [27].

